# Breaking Through Biology’s Data Wall: Expanding the Known Tree of Life by Over 10x using a Global Biodiscovery Pipeline

**DOI:** 10.1101/2025.06.11.658620

**Authors:** Oliver Vince, Phoebe Oldach, Valerio Pereno, Marcus H Y Leung, Carla Greco, Gus Minto-Cowcher, Saif Ur-Rehman, Keith Y K Kam, William Chow, Emma Bolton, Bupe R Mwambingu, Nadine Greenhalgh, Ineke Knot, Leif Christoffersen, Marlon Clark, Robert Pecoraro, Aaron W Kollasch, Tanggis Bohnuud, Matthew Bakalar, Philipp Lorenz, Glen Gowers

## Abstract

Advancements in the life sciences have always been built upon our collective understanding of life on Earth. Now, the rise of generative biology - the use of AI foundation models to design, generate, and annotate proteins, pathways and therapeutics - is creating unprecedented demand for large, diverse biological sequence datasets. While a limited subset of such data can be generated in clinical or laboratory settings, the vast majority of the training data for unsupervised models must be sourced from the natural world - the product of nearly four billion years of evolutionary history.

However, the public databases that currently supply this data, while foundational to research, were established to aggregate results from academic experiments, not as training datasets for machine learning. Their human-centric data structure limits model performance due to redundancy, taxonomic and geographic bias, limited biological context, and inconsistent provenance. With 68% of all sequence data in the SRA database coming from just 5 species, this is one of the most severe class imbalance problems ever encountered in AI. Legal and infrastructural constraints further exacerbate this bottleneck.

To address these limitations and support scalable model training, we introduce BaseData: the largest and fastest-growing biological sequence database ever built, and the first purpose-built for training foundation models. As of late 2024, BaseData contained 9.8 billion novel genes, representing more than a 10-fold expansion in known protein diversity after accounting for redundancy. BaseData also contains more than 1 million species not represented in other genomic databases. Its partnership-driven data supply chain across 26 countries and autonomous regions enables growth of up to 2 billion novel genes per month, far exceeding public repositories. All data is collected under benefit sharing agreements using standardized protocols and structured using graph-based, ontology-rich metadata that preserves evolutionary context.

BaseData represents a new, ethically-grounded infrastructure for training biological foundation models, complementing public efforts and enabling the next era of generative biology.

**Short Abstract:** Progress of AI in biology is now being limited by the availability of high-quality biological sequence data from nature. To get past this data wall, we introduce BaseData™, a new biological database built on top of a global, scalable biodiscovery pipeline. As of 2024, BaseData had already expanded the known protein universe by over 10x.

## 1. Introduction: The Data Wall in Biology

Humanity’s progress has always been intimately tied to its ability to understand and harness the natural world. From the agricultural revolution and Aristotle’s Historia Animalium (Aristotle and Thompson 1910) to deciphering the genetic code and the engineering of synthetic genomes, our expanding biological knowledge has revealed the inner workings of life and enabled us to reshape it to deliver curative medicines, resilient crops and sustainable materials.

Now, as our ability to digitise biology advances (Bunne et al. 2024; Hey 2012; King et al. 2009) alongside exponential gains in computational power (Garisto 2024) and algorithmic efficiency (Krichen and Abdalzaher 2024), we stand on the cusp of a new era driven by foundation models - large-scale machine learning systems capable of capturing generalisable representations across entire biological domains.

In this paper, we explore biological foundation models as a transformative new approach for understanding and manipulating biology. We then present our hypothesis that the emerging data bottleneck is leading to an observable plateau in model performance (Notin 2025) and constitutes a limit to further progress in the field. We then introduce BaseData, the largest and fastest-growing biological sequence database designed specifically for AI training, and demonstrate how it offers a scalable, inclusive, and purpose-built foundation for training the next generation of biological AI models.

We continue by analysing the structural limitations of existing public sequence databases and identify the key reasons behind their poor alignment with the needs of modern machine learning. Finally, we present the data supply chain that BaseData is built upon: Basecamp Research’s novel, partnership-based framework for large-scale, ethically-sourced data collection from biodiversity.

### 1.1 The Promise of Biological Foundation Models: The Next Leap in Understanding Life on Earth

Artificial intelligence models trained on massive biological datasets are enabling unprecedented insights into biological structure, function, and design, ushering in a transformative new era in life sciences. These models compress the collective evolutionary record into a compact, reusable form of biological knowledge. Akin to an experienced scientist drawing on years of accumulated knowledge, these models can, when pre-trained at scale, efficiently infer complex structure-function relationships, regulatory codes, and fitness landscapes. This enables them to make accurate predictions in unfamiliar biological settings with minimal input, significantly reducing the reliance on large-scale experimental screening (Zhang et al. 2024; Puniya 2025).

Foundation models have already transformed multiple scientific and technical frontiers. In natural language processing, models such as GPT (“Introducing GPT-4.5” 2025) and PaLM (Narang and Chowdhery 2022) demonstrated that the implicit rules of complex systems can be learned from unlabelled data at scale, enabling generalisation across diverse tasks with minimal supervision. Similar strategies are now advancing nuclear fusion research (Churchill 2025), where machine learning is used to model turbulent plasma behaviour and optimise reactor control systems, and astrophysics, where models trained on petabytes of telescope and simulation data are helping to unravel galaxy formation and dark matter distributions (Shukla, Bandyopadhyay, and Karelia 2025; M. Huertas-Company and Lanusse 2023). These domains share key challenges with biology: extreme complexity, sparse mechanistic understanding, and a dependence on large, heterogeneous datasets - making biology a natural and fertile ground for the next generation of foundation models.

The ’coding languages’ of DNA, RNA, and proteins are universally shared across biology and exhibit an inherent regularity, making them an especially strong foundation for training biological foundation models. Such models have already been shown to excel in annotation, prediction, and generative design tasks with notable examples including Google Deepmind’s AlphaFold2 (Jumper et al. 2021; Abramson et al. 2024), EvolutionaryScale’s ESM series (Hayes et al. 2025), Profluent’s ProGen series (Bhatnagar et al. 2025), The Arc Institute’s Evo series (Nguyen, Poli, Durrant, Kang, et al. 2024; Brixi et al. 2025) and Tatta Bio’s gLM series (Hwang et al. 2024; Cornman et al. 2024), and InstaDeep’s Nucleotide Transformer (Dalla-Torre et al. 2025): more than 360 biological foundation models have now been published to date (Maug 2024).

Just as human language models can generalise across dialects and topics, biological foundation models have the potential to integrate information across genes, proteins, cells, organisms, and environments. The generalisability of these models is key for large scale adoption across R&D workflows: it opens the door to models that could theoretically be capable of predicting behaviors or designing complex, multi-component biological systems such as synthetic consortia, personalised therapeutic regimes, or *de novo* genomes with levels of speed, sophistication and precision that far exceed what evolution or human engineering has so far produced and likely will ever be able to produce.

This generalisability can also inform accurate predictions in unfamiliar biological settings with minimal input, significantly reducing the reliance on large-scale experimental screening. This reduction in experimental burden unlocks the possibility of repurposing this experimental capacity away from generating large pre-training datasets to generating smaller but more relevant pre-clinical or clinical datasets for fine-tuning.

The potential of small, high-fidelity datasets to align the output of protein language models with protein design objectives was first demonstrated in a seminal paper by the Church lab (Biswas et al. 2021). Using a strategy of iterative low-N experimental assays with as few as 24 mutants per round, the authors trained a top model on eUniRep to design improved variants of green fluorescent protein (GFP) with fluorescence output on par with the highly-engineered Superfolder GFP. Another compelling demonstration is EVOLVEpro (Jiang et al. 2025), a framework that integrates the ESM-2 protein language model with few-shot active learning to optimize protein function. In evaluations across diverse protein families, including T7 RNA polymerase, a miniature CRISPR nuclease, a prime editor, an integrase, and a monoclonal antibody, EVOLVEpro identified high-performance variants using as few as 10–16 experimental data points per round, achieving up to 100-fold improvements in desired properties over five rounds. These strategies enable biological foundation models to adapt general biological patterns to specific tasks, akin to an expert biologist forming hypotheses after limited preliminary data. As foundation models continue to advance, such low-N design workflows hold the potential to minimize or even obviate the need for extensive high-throughput screening. Such an approach would accelerate the traditional development pipeline, enabling rapid, low-cost, and highly personalised medicine, where candidate molecules or interventions are virtually optimised before they ever reach the lab bench.

Another powerful aspect of these models is their growing ability to interact with natural language interfaces (Fallahpour et al. 2025; Ghareeb et al. 2025). Human scientists are adept at formulating goals and problems in language, but limited in their ability to convert these intuitions into actionable biological designs. Foundation models act as powerful intermediaries: systems capable of translating human intent, expressed in plain language, into executable biological solutions.

In summary, biological foundation models mark the beginning of an important shift towards understanding and engineering biological systems in their full complexity. By compressing evolutionary history into a generalisable and reusable scaffold, these models can guide design with minimal experimental data, redefining the boundaries of feasibility in synthetic biology, medicine, and environmental engineering. However, like their counterparts in language and vision, the performance of these models is tightly linked to the scale and diversity of their training data. As models advance, they are fueling demand for new, diverse and well-contextualised biological data that vastly exceeds the capacity of current data infrastructures.

### 1.2 The Data Wall Holding Back Foundation Models in Biology

Biological foundation models promise to transform our ability to understand, program, and engineer biological systems. Early models with large-scale training on biological sequences have demonstrated impressive generalisation, but, in contrast to fields such as language, vision, and chemistry, more recent developments show that model growth is slowing down (**Figure 1E)**. An analysis of over 360 biological foundation models found a slowdown in development beginning around 2021, just as many models reached or exceeded the training dataset size of UniRef50, the most commonly used protein training dataset (Maug 2024; Villalobos 2025). This inflection point suggests that the primary constraint may no longer be algorithmic innovation, but the availability of sufficiently diverse and well-curated biological data. As these models continue to scale, their performance increasingly hinges on overcoming the limitations of existing public sequence repositories.

**Figure 1:**
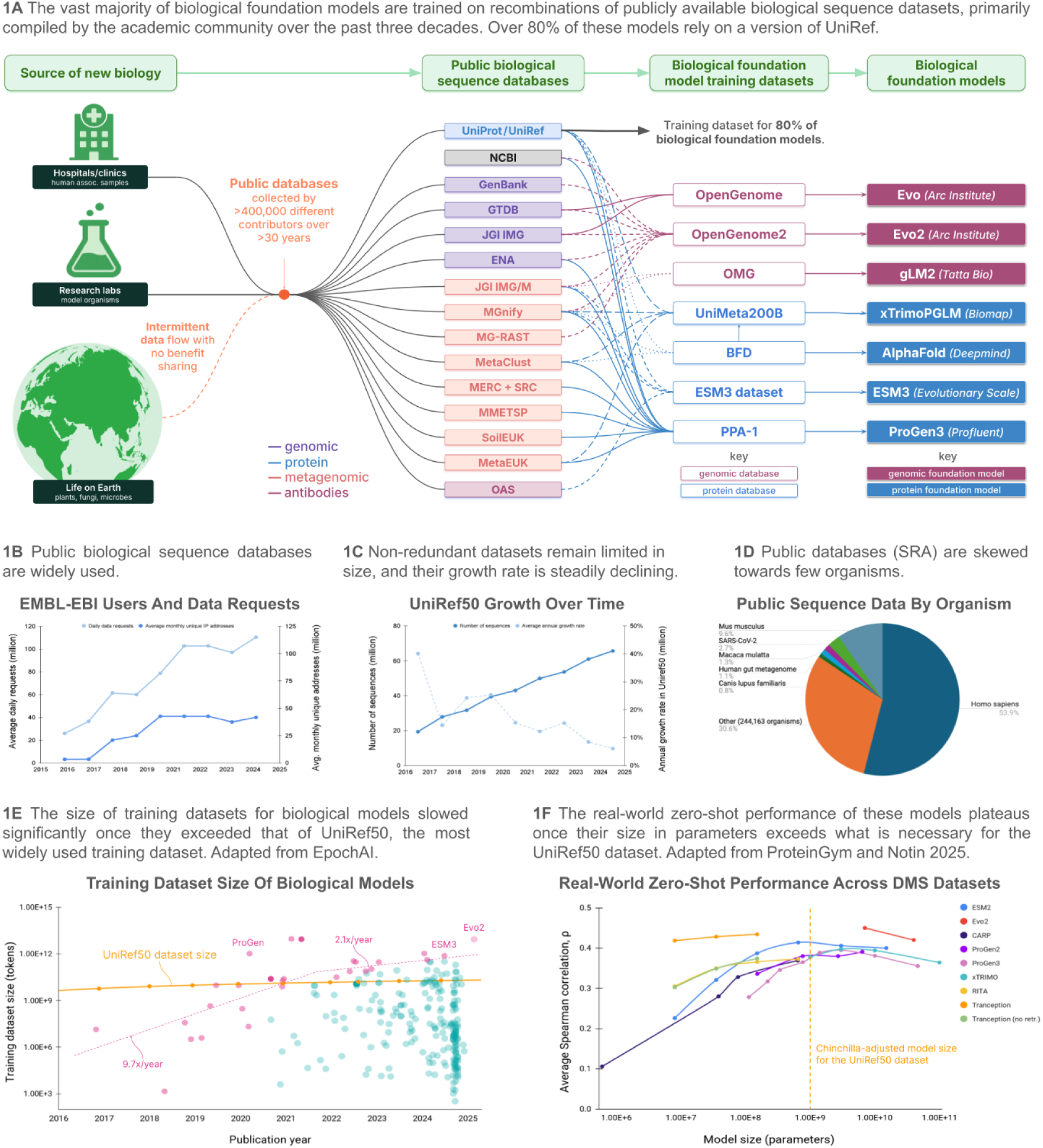
The data wall and performance plateau for foundation models in biology

#### All Biological Foundation Models Train on the Same Limited Public Datasets

In contrast to specialised AI models designed for narrow biological tasks, foundation models aim to acquire a generalisable understanding of biological systems. Access to training datasets that comprehensively reflect the full breadth and complexity of biological sequence space, across all domains of life, ecological niches, and molecular functions are needed to achieve this generalisation.

In practice, such sequence data originates from three main sources: clinical sequencing (e.g., human genomics and microbiomes), laboratory experiments (e.g., model organisms, synthetic constructs), and biodiversity sampling (e.g., environmental and metagenomic sequencing). Researchers collecting sequence data from clinical, laboratory, or environmental sources commonly deposit data in publicly available biological sequence repositories, such as the European Nucleotide Archive (ENA) (Leinonen, Akhtar, et al. 2011) and the Sequence Read Archive (SRA) (Leinonen, Sugawara, et al. 2011), which represent the mainstay of the global exchange of nucleotide sequence data.

These databases are widely used: EMBL-EBI (the research institute that operates and maintains ENA) reported over 115 million data requests per day from more than 40 million users annually in 2024 (Beagrie and Houghton 2021) **(Figure 1B)**. Additionally, as an illustrative example of the commercial dependence on these public databases, analysis of 4 major patent databases (USPTO, EPO, JPO, KIPO) by the authors found that 66% of patents have at least one claimed sequence with over 90% sequence ID to a sequence on RefSeq.

The curated databases that stem from these public repositories such as UniRef (Suzek et al. 2015), GenBank (Benson et al. 2013), RefSeq (Goldfarb et al. 2025) and MGnify (Richardson et al. 2023) constitute the most comprehensive and high-quality sources of biological sequence data available. Open access to these databases stands in contrast to large clinical genomics databases and proprietary laboratory datasets within the life sciences industry, which are typically restricted and therefore unavailable for training foundation models.

Nearly all published foundation models today that incorporate biological sequence data into their training source their pretraining data from these free, publicly accessible online sequence repositories (**Figure 1A)**. More than 80% of foundation models are trained on UniRef, one of the largest and most curated public protein sequence databases (Maug 2024). This includes protein structure prediction models (e.g., AlphaFold2 (Jumper et al. 2021), ESMFold (Lin et al. 2023), OpenFold (Ahdritz et al. 2024), and RoseTTaFold (Baek et al. 2021), protein-ligand models (e.g., NeuralPlexer (Qiao et al. 2024), protein function models (e.g., CLEAN (Yu et al. 2023), ProteInfer (Sanderson et al. 2023), and ProteinVec (Hamamsy et al. 2023) and protein generation models (e.g., Chroma (Ingraham et al. 2023), ProteinGAN (Repecka et al. 2021), ProGen (Madani et al. 2023), ProtGPT2 (Ferruz, Schmidt, and Höcker 2022), and EvoDiff (Alamdari et al. 2023).

#### Small, Skewed and Stagnant: Public Sequence Databases Are a Poor Foundation for AI Training

Despite now being central to training biological foundation models, public sequence databases such as UniRef were never intended for machine learning applications but rather to consolidate the outputs of diverse academic experiments from around the globe, prioritising human interpretability, reproducibility, and data sharing. As a result, foundation models trained on these datasets inherit several structural limitations: modest effective size and information content, taxonomic and ecological biases, limited contextual metadata, inconsistent curation standards, and slow growth in non-redundant content.

At the raw data level, the Sequence Read Archive (SRA) is vast, comprising over 50 petabases of sequencing data from more than 27 million datasets as of the end of 2023 (Chikhi et al. 2024). However, analysis by the authors found that more than half (53.9%) of all sequencing data in the SRA is *Homo sapiens*, followed by 9.6% from *Mus musculus* and 2.7% from *SARS-CoV-2*. In total, 68.3% of all SRA sequence data comes from just 5 species (**Figure 1D)**. Geographically, 70% of all SRA submissions originate from only ten countries (**Figure 4D)**. These imbalances reflect historical research priorities, but they now severely limit the capacity of foundation models to generalise across the full diversity of life.

Fewer than 2.3 million species have been formally described (“Catalogue of Life” 2025), yet estimates suggest there may be over 8–15 million eukaryotic species and more than one trillion microbial taxa globally (Locey and Lennon 2016). As of 2025, fewer than 22,000 out of a target of 1.8 million eukaryotic species had been sequenced to high-quality reference standards (“Earth Biogenome Project” 2025). Even among described species, representation is uneven, with entire branches of the tree of life, for example, fungi, protists, deep-sea microbes, and insects, severely underrepresented or absent.

Crucially, very little of this raw data ultimately reaches the training corpora used in machine learning. From the SRA, sequence data must pass through numerous intermediate steps, including assembly, gene prediction, quality filtering, clustering, and annotation, before entering curated resources like UniProt (UniProt Consortium 2023; Chikhi et al. 2024). As a result, the vast majority of original sequence information is lost or excluded along the way.

UniRef50, the clustered and non-redundant version of UniProt most commonly used in protein language model training, contains fewer than 70 million sequences, representing a loss of more than 99.999% of the raw sequence content when compared to the 50 petabases in the SRA. For the small number of curated sequences that result they often still lack contextual metadata, including functional annotations, taxonomic origins, and provenance labels.

Despite exponential reductions in sequencing costs and rapid growth in raw data generation, the annual growth rate of UniRef50 has steadily declined, now below 10% per year **(Figure 1C)**. This growing mismatch underscores a systemic bottleneck: the scale and quality of training datasets remain bounded by the constraints of legacy infrastructures and fragmented voluntary data deposition.

#### Mining for More: Other Attempts to Expand the Dataset Frontier

As the field has moved towards larger foundation models, a wider range of publicly available sequence data has been mined (**Figure 2A)**. These efforts have collectively raised the total number of biological sequences suitable for model training to trillions of genomic tokens (nucleotides) and billions of protein sequences **(Figure 2B and 2C)**.

**Figure 2:**
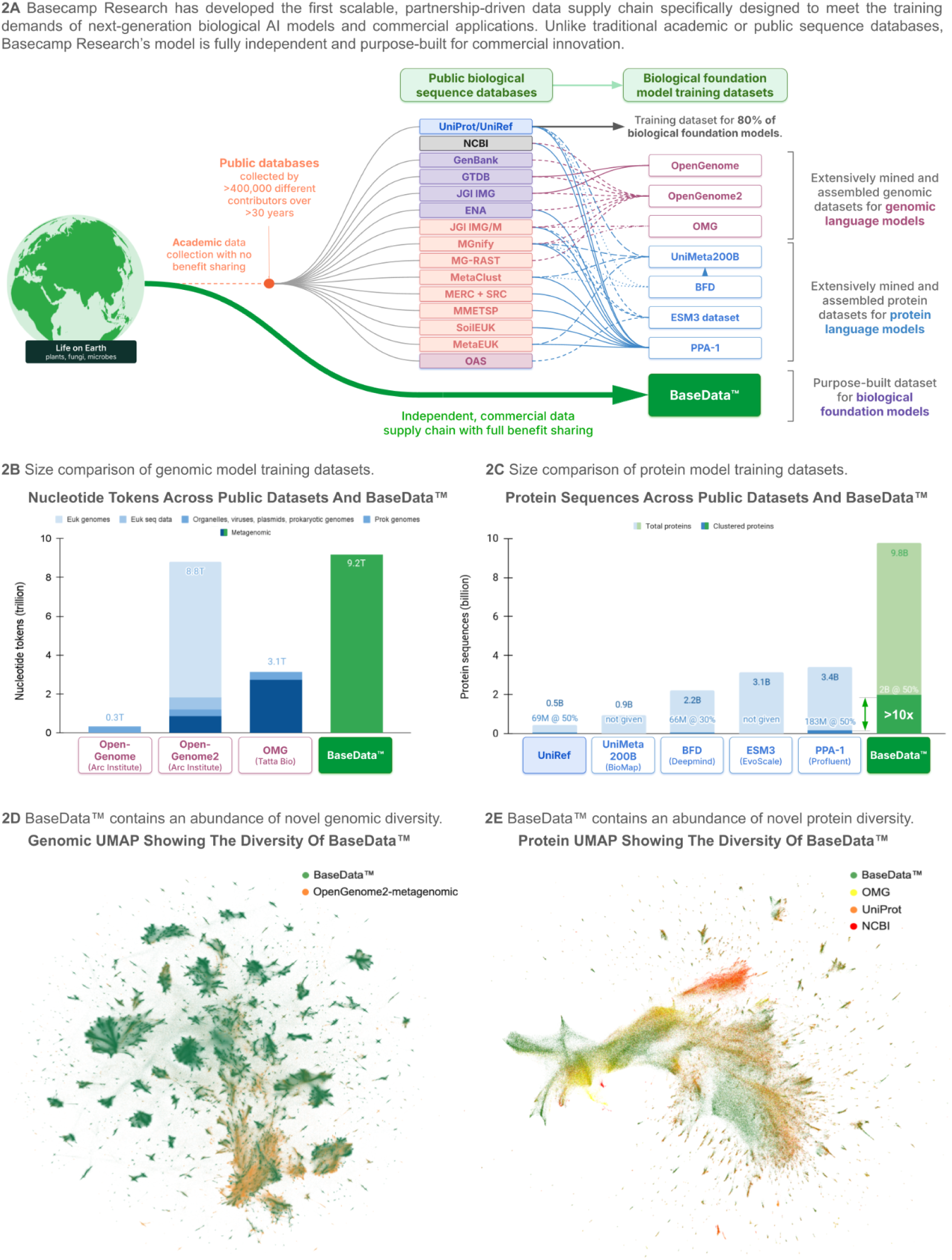
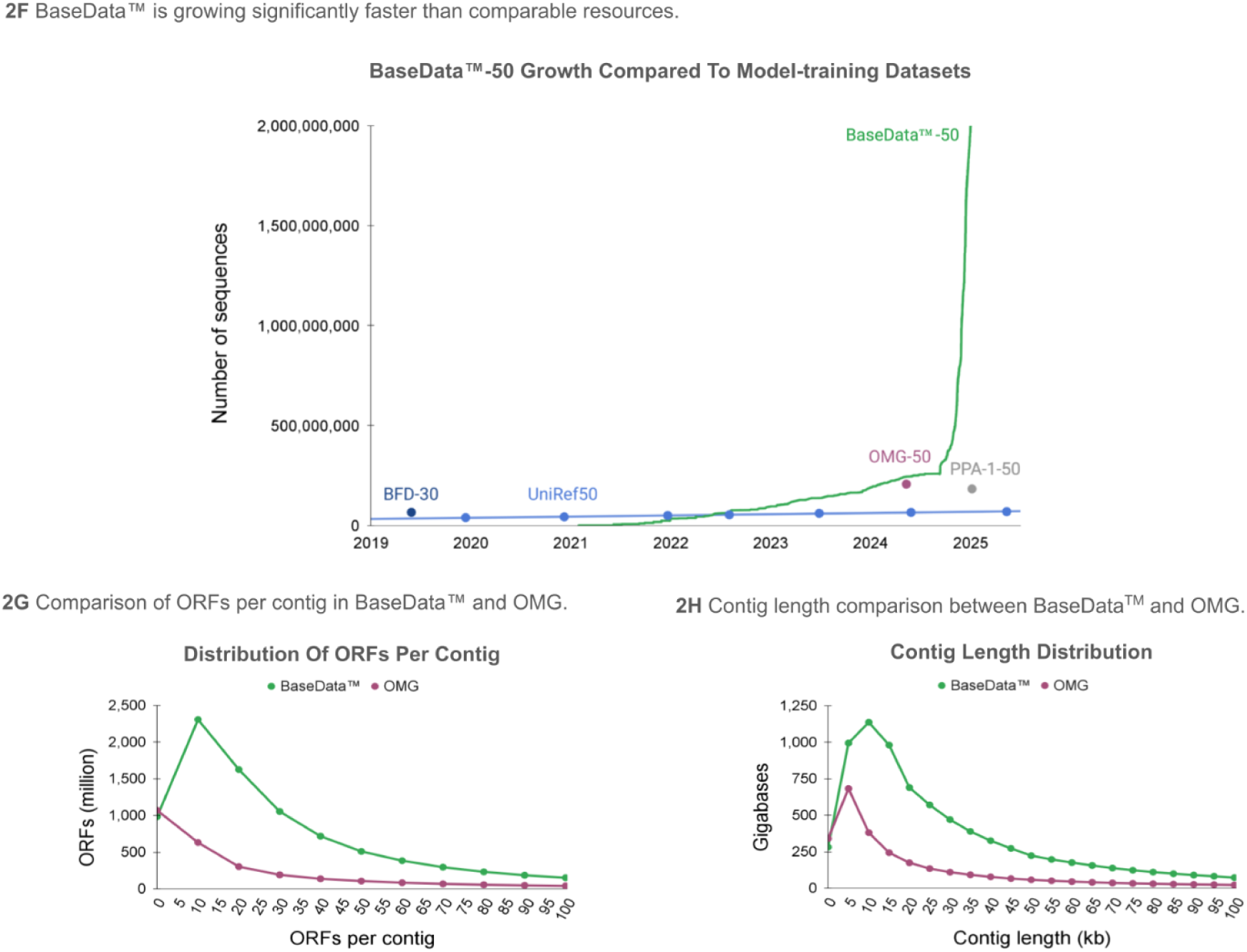
Introducing BaseData™ - a new, independent and fully-scaled database purpose-built for AI model training

For genomic foundation models, these resulting databases include OpenGenome and OpenGenome 2 (for training the Evo (Nguyen, Poli, Durrant, Thomas, et al. 2024) and Evo2 (Brixi et al. 2025) models, respectively) and the OMG dataset (for training the gLM2 model (Cornman et al. 2024). At 8.8 trillion genomic tokens (nucleotides) OpenGenome2 is the largest published dataset ever curated for genomic language model training with the resulting model, Evo2, at 40 billion parameters, the largest genomic model to date. However, less than 10% of the tokens (854 billion) in the OpenGenome2 dataset come from non-redundant metagenomic data, with 80% of the tokens (7 trillion) coming from eukaryotic genomes, which makes up only a small fraction of the overall tree of life **(Figure 2B)** (Larsen et al. 2017). The largest published metagenomic dataset assembled for genomic model training is the OMG dataset. With 3.1 trillion genomic tokens in total, 87% of these (2.7 trillion) come from non-redundant metagenomic data, with the remainder coming from prokaryotic genomes **(Figure 2B).**

For protein foundation models, xTrimoPGLM (Chen et al. 2023) was trained on the UniMeta200B dataset, which incorporates data from BFD (Steinegger 2019), a 2.2 billion sequence dataset originally assembled by Google DeepMind and the Technical University of Munich for AlphaFold2 (Jumper et al. 2021) and AlphaFold3 (Abramson et al. 2024). ESM3 was trained on the ESM3 dataset (3.1 billion sequences), and, at the largest size, the ProGen3 model was trained on the PPA-1 dataset (3.4 billion sequences) **(Figure 2B)**.

However, these datasets are not independent; they reflect different combinations, filtering strategies, and deduplication thresholds applied to the same foundational sources, such as UniProt, JGI IMG, and MGnify **(Figure 1A)**. As a result, they all inherit the same structural limitations: high redundancy, uneven taxonomic and geographic representation **(Figure 1D and 4D)**, limited contextual metadata, and relatively slow growth in non-redundant content **(Figure 1C)**.

These limitations become evident when removing redundancy - a standard practice in AI training workflows to prevent models from overfitting to highly represented regions of sequence space. UniRef loses 84.8% when clustered at 50% (454 million down to 69 million sequences), BFD loses 97% when clustered at 30% (2.2 billion down to 66 million sequences), the PPA-1 loses 95.6% when clustered at 50% (3.4 billion down to 183 million sequences), and OMG loses 93.7% when clustered at 50% (3.3 billion down to 207 million) (Bhatnagar et al. 2025) **(Figure 2C).**

Once redundancy is accounted for, even the largest biological sequence curation initiatives have expanded the known protein diversity by less than fourfold (Bhatnagar et al. 2025). By the standards of other AI domains, this represents an unusually constrained training corpus.

#### Additional Structural Barriers to Machine Learning

Beyond size and bias, public biological sequence datasets face several compounding challenges. Annotation depth is limited (less than 1% of UniProtKB entries are manually curated (“UniProt” 2025) and many sequences lack functional labels or reliable taxonomic assignment. Short fragments dominate metagenomic data, offering little structural or contextual signal for machine learning. Provenance is often unclear, with wide variability in experimental methods and bioinformatic pipelines.

Databases also remain siloed, with inconsistent taxonomies and poor cross-referencing between resources. These limitations, combined with the legal, regulatory and reputational issues discussed elsewhere in this paper, undermine the construction of integrated, high-quality training sets and restrict models’ ability to learn biologically meaningful representations at scale.

### 1.3 Hitting the Data Wall: Model Growth is Slowing Down

The growth trajectory of biological foundation models is encountering a significant inflection point: the availability of high-quality, diverse training data is becoming a limiting factor. This slow down was first quantified by the team at EpochAI who analysed more than 360 biological AI models and observed *“rapid scaling from 2017-2021, followed by a notable slowdown in biological model development”* (Villalobos 2025). **Figure 1E** illustrates this trend and has been adapted directly from the EpochAI analysis.

To assess the extent to which training data scale, quality, and diversity may be contributing to the observed slowdown in biological model scaling, we plotted the size of UniRef50 (the most widely used non-redundant dataset for protein language model training) in amino acid tokens alongside model training dataset sizes in **Figure 1E**.

This shows that the observed slowdown in model growth begins as model training datasets start to exceed the size of UniRef50. While this constitutes a correlation rather than a definitive causal link, the timing is notable. Additionally, even the largest curated datasets used in model training, such as PPA-1 (Bhatnagar et al. 2025), BFD (Jumper et al. 2021), and OMG (Cornman et al. 2024), expand the known protein sequence diversity by less than fourfold once redundancy is removed **(Figure 2C)**. Taken together, these trends suggest that biological foundation models are now outpacing the availability of high-quality public data.

This pattern mirrors phenomena observed in other AI domains. In natural language processing (NLP), early advancements were propelled by scaling model sizes and training on expansive, high-quality text corpora (Kaplan et al. 2020). However, as models surpassed billions of parameters, the scarcity of diverse, high-quality text data became apparent, leading to reliance on broader, noisier sources. Similarly, in computer vision, initial breakthroughs were driven by datasets like ImageNet (Krizhevsky, Sutskever, and Hinton 2017). Yet, performance gains plateaued until the introduction of much larger datasets such as JFT-300M and LAION-5B, which offered greater visual diversity and scale (Sun et al. 2017; Schuhmann et al. 2022).

Biological foundation models, particularly those modeling proteins and genomic sequences, are now facing a similar data bottleneck. Unlike NLP or vision models that can tap into ever-expanding sources of text and imagery, most biological foundation models still depend heavily on the comparably small and slow growing public sequence databases **(Figure 1A)**.

### 1.4 From a Data Wall to a Performance Plateau

The slowdown in dataset growth is now being mirrored by a corresponding plateau in model performance. One of the clearest illustrations of this plateau can be seen in the result of the comprehensive ProteinGym benchmarks, which evaluated more than 70 high-performing protein language models across more than 250 deep mutational scanning assays (Notin et al. 2023). Each model was evaluated in a zero-shot setting, reflecting real-world scenarios where predictions must be made without task-specific training or abundant experimental data. ProteinGym is particularly valuable for its breadth and its emphasis on biologically meaningful mutations that directly impact protein function and disease relevance.

The aggregated results in ProteinGym reveal a clear saturation point: once models surpass a certain parameter threshold, additional scaling yields little improvement in predictive accuracy, and in some cases, leads to decreasing zero-shot performance **(Figure 1F)** (Notin 2025).

Again, to assess the extent to which data limitations may be contributing to the observed performance plateau in biological foundation models, we plotted a vertical line on **Figure 1F** representing the model size predicted by the Chinchilla scaling ratio for a dataset the size of UniRef50 (Hoffmann et al. 2022). Originally developed in the context of natural language processing, the Chinchilla model demonstrated that optimal performance is achieved by an optimal ratio of carefully balancing model size and dataset scale: for a fixed data budget, smaller models trained longer typically outperform larger models trained on the same amount of data. While this principle was derived from autoregressive language modeling, it provides a useful heuristic for thinking about trade-offs in other domains. Beyond ProteinGym, performance plateaus have also been observed where larger model sizes no longer yield proportional gains (Weissenow and Rost 2025; Nijkamp et al. 2023; Hesslow et al. 2022).

Together, these observations are consistent with the conclusion that the field has entered a data-constrained scaling regime and are consistent with the findings of (Ding and Steinhardt 2024) that show that protein language models inherit strong species-level biases from their training data, disproportionately favoring taxa that are overrepresented in public sequence databases (Ding and Steinhardt 2024). These biases limit generalisability and degrade performance on underrepresented or evolutionarily distant species.

It is therefore reasonable to assume that without access to substantially more diverse, well-annotated, and evolutionarily representative training datasets, further increases in model size are unlikely to yield meaningful improvements in zero-shot performance or real-world biological applications.

It is important to note that this correlation between the predicted scaling threshold and the observed flattening of performance is not definitive evidence of causality. It has been argued that even models trained on larger and more diverse biological sequence datasets may not achieve perfect prediction of the fitness landscape, due to confounding factors such as phylogenetic bias and evolutionary constraint. Weinstein et al. (2022) reinforce this point, showing that the relationship between extant sequences and fitness is shaped by deep evolutionary histories, making it difficult for models to disentangle causality from correlation (Weinstein et al. 2022). In this light, even a more diverse dataset may be insufficient if its structure reflects the same inherited biases.

#### Innovating Around the Wall: Other Strategies to Overcome Data Limitations

The emergence of a data-constrained scaling ceiling has prompted a range of efforts aimed at extending model capabilities in the face of limited training data. Several recent studies have explored architectural, algorithmic, and data-centric strategies to mitigate these constraints.

ProGen3 demonstrated that aligning large foundation models with targeted experimental datasets such as mutational fitness or protein stability can significantly improve performance on practical design tasks, including out-of-distribution prediction and *in vitro* expression (Bhatnagar et al. 2025). Additionally, test-time scaling approaches that increase computational resources during inference, allowing models to generate more refined outputs without additional training have shown promise (Nabla Bio 2025). Meanwhile, Fournier et al. argue that increasing model scale does not guarantee better biological performance, particularly when training data fails to accurately represent the underlying fitness landscape. Instead, they advocate for improved model architectures, improved data quality, and rigorous evaluation as more reliable paths toward improved performance (Fournier et al. 2024).

Nonetheless, the strategies emerging across academia and industry, ranging from dataset diversification and targeted alignment to test-time scaling and architectural refinement, demonstrate that meaningful improvement is still possible. What remains uncertain, and increasingly urgent, is the extent to which performance ceilings can be lifted by improved datasets alone. Systematic construction of high-quality, biologically grounded, and scaled datasets will be essential to answer this question.

## 2. BaseData: The Largest And Fastest Growing Biological Sequence Database

### 2.1 A Fully Independent and Purpose-Built Data Supply Chain

BaseData was created to break through the data wall that we predict to be limiting the performance of foundational biological sequence models. It is the first biological sequence resource built specifically for training foundation models: independent, continually growing, maximally diverse, and globally sourced with full legal and regulatory clarity (**Figure 2A**). Unlike existing public databases, BaseData is growing rapidly and is built to continue to scale in both size and diversity. It captures deeper biological signals, includes longer and more informative sequences, and offers a significantly broader view of the tree of life. Every sequence is fully traceable and collected with commercial use in mind, enabling fast and compliant integration into industrial and clinical pipelines. This sets a credible and scalable foundation for training the next generation of biological foundation models.

Public biological sequence databases have been built over decades through the aggregated contributions of hundreds of thousands of researchers, institutions, and sequencing projects. While this aggregative model has driven foundational progress in biology, it is inherently slow, uneven, and difficult to steer.

BaseData takes a fundamentally different approach. Built on a fully independent, purpose-engineered, and scalable data supply chain, it sources biological diversity through structured commercial partnerships founded on equitable bilateral access and benefit sharing agreements across more than 120 field sites in 26 countries and autonomous regions.

### 2.2 Scale: The Largest Biological Sequence Database

BaseData currently represents the largest biological sequence database ever constructed for AI model training in both genomic and protein space. Its total nucleotide content at the end of 2024 exceeded 9.2T tokens, surpassing the 8.8T in OpenGenome2, and almost 3x larger than OMG, at 3.1T tokens (**Figure 2B**). When considering the more directly comparable metagenomic portions of the OpenGenome2 and OMG datasets, BaseData is over 10x larger than OpenGenome2 and over 3x larger than OMG (**Figure 2B**).

In the protein domain, BaseData also sets a new benchmark. BaseData contains 9.8 billion highly filtered and curated protein sequences, all coming from genomic contigs that are at least 2kb long (**Figure 2C**). In absolute size, this makes BaseData 21.5x larger than UniRef, 10.4x larger than UniMeta200B, 4.4x larger than BFD, 3.1x larger than the database used to train ESM3 and 2.9x larger than PPA-1, the database used to train ProGen3 (**Figure 2C**).

BaseData leads the field not just in sheer size but in protein diversity. BaseData-50 (BaseData clustered at 50% sequence ID) is 31.9x larger than UniRef50 and 12x larger than the PPA-1, also clustered at 50%. As the PPA-1 represents the largest protein database yet curated for foundation models in biology, this means that BaseData can be considered to have expanded diversity of proteins known to science by over an order of magnitude.

### 2.3 Velocity: The Fastest Growing Biological Sequence Database

While BaseData has already demonstrated orders-of-magnitude expansions on biological sequence knowledge, importantly, its scale is not static. BaseData is actively growing, purpose-built to meet the growing data demands of models at the multi-billion- and trillion-parameter level. In 2024 BaseData-50 grew by over 10x in a year, with a maximum monthly sequence growth rate of 2 billion novel, high-quality protein sequences per month, a pace that far exceeds that of comparable clustered datasets (**Figure 2F**). Crucially, this growth of 50% clusters reflects the discovery of new biological diversity, not the repeated sampling of known sequences.

### 2.4 Novelty: Capturing New Biological Diversity and New Species

In addition to its scale, what sets BaseData apart is its ability to capture previously unobserved regions of biological sequence space. A UMAP projection of genomic sequences reveals that BaseData occupies large areas of sequence space absent from the metagenomic portion of OpenGenome2 (**Figure 2D**). Similarly, a UMAP projection of clustered protein sequences reveals that BaseData occupies large areas of sequence space absent from widely used protein datasets such as UniProt, NCBI, and OMG_prot50 (**Figure 2E**). This novelty extends beyond sequence space into taxonomic space: BaseData includes over 1 million new species, as defined by unique Operational Taxonomic Units, not found in GTDB or OMG, highlighting its unprecedented contribution to species-level discovery (**Figure 3E**). An example of the diversity of marker gene RecA in BaseData is shown in the phylogenetic tree in **Figure 3F**. This is most likely an under-estimate given that not all ground-truth marker genes were present in the contigs and bins post assembly. Analyses across diverse commercially-relevant protein families such as recombinases, hydrolases, and ATP synthases demonstrate that BaseData consistently captures more sequence-level and phylogenetic diversity than any existing dataset, finding more potential starting points for biological functions important for the development of new therapeutics and industrial solutions **(Figure 3A–D)**.

**Figure 3:**
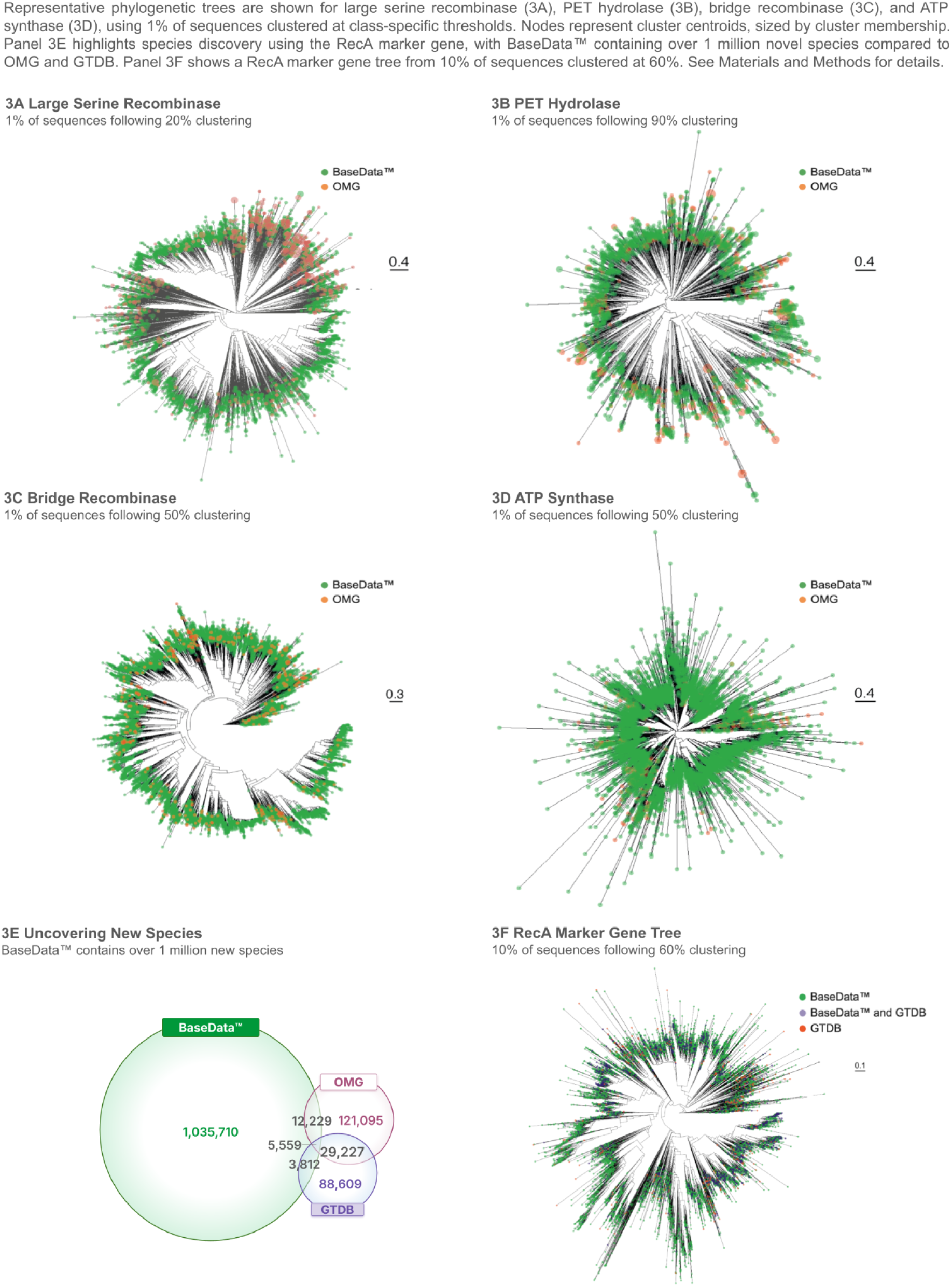
BaseData™ contains higher diversity than comparable resources

**Figure 4:**
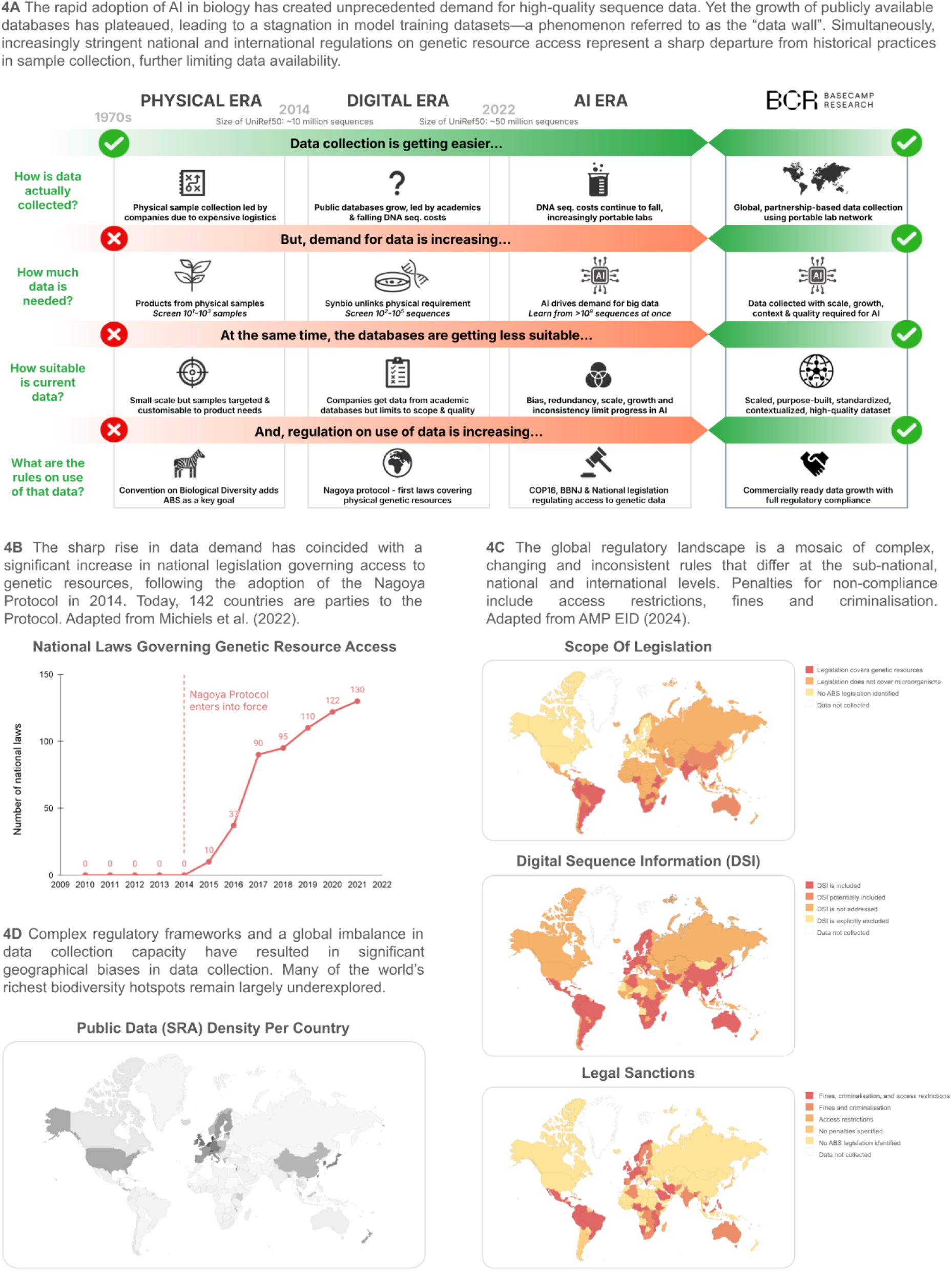
Why is there a data wall in biology?

### 2.5 Context: Richer Sequences, More Biological Signal

Beyond its scale and diversity, BaseData offers a step-change in the biological richness of the data captured. Thanks to purpose-designed extractions and sequencing modalities, assembled sequences in BaseData exhibit significantly longer assemblies, with 6.77 trillion nucleotides appearing in contigs longer than 10kb, compared to only 1.93 trillion for OMG (**Figure 2H**) and contain more open reading frames (ORFs) per contig compared to leading public metagenomic datasets such as OMG (**Figure 2G**). This enables a higher likelihood of capturing complete biological mechanisms within individual records.

Each sequence within BaseData is also embedded within a deep metadata layer capturing environmental, chemical, and physical parameters, as well as genomic and metagenomic context. **(Figure 5B).** This enables models to learn from molecular patterns and interconnected biological systems, including co-occurrence of genes, horizontal gene transfer networks, and ecological associations.

**Figure 5:**
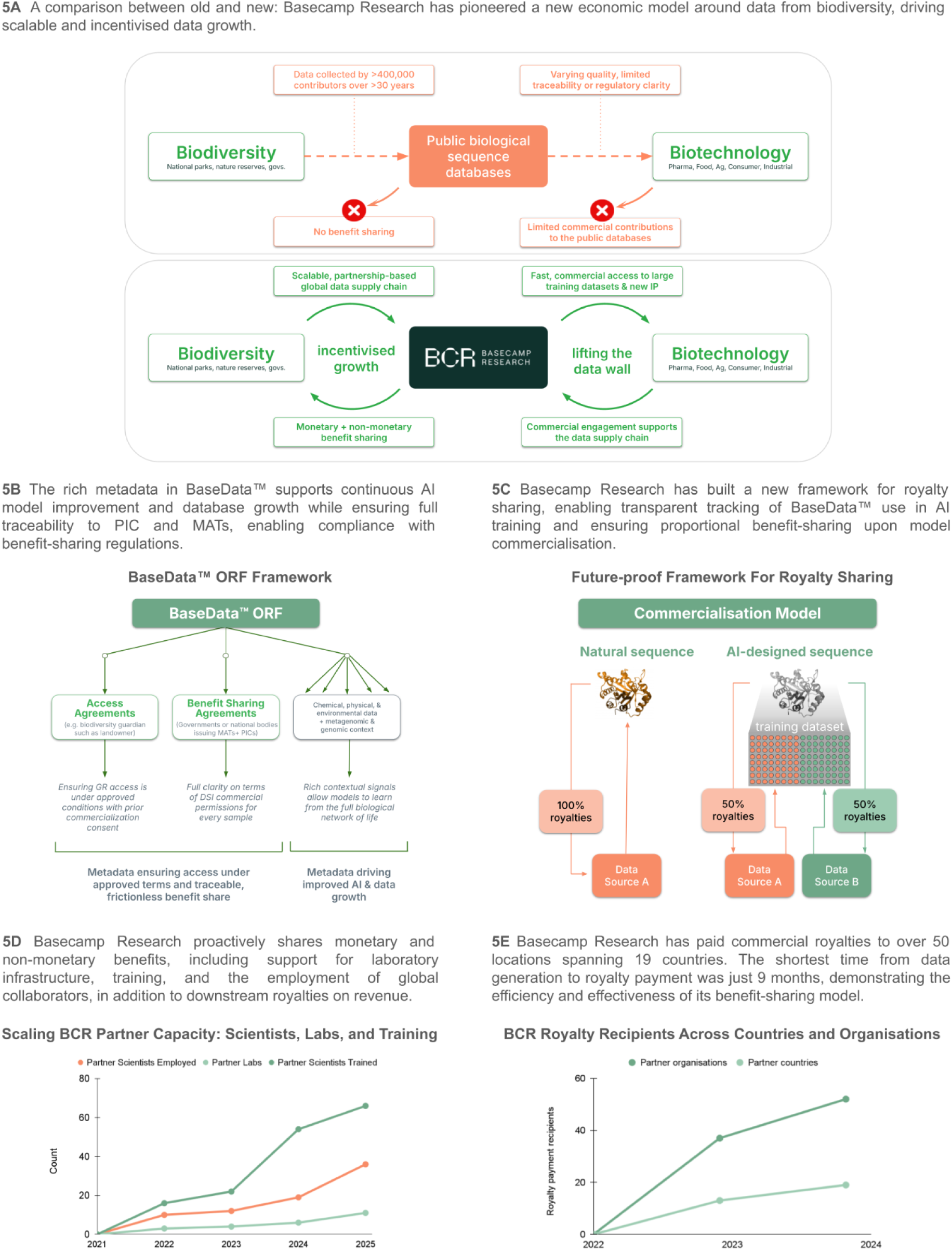
Basecamp Research’s data supply chain - a new, partnership-based and incentivised approach to rapidly expand our knowledge of life on Earth

## 3. Understanding the Data Wall: Why Do We Know So Little About Life on Earth?

### 3.1 A Demand Shock: AI Models Have Outgrown Public Biological Data

The emergence of biological foundation models marks a sharp inflection point in how biological sequence data is used (**Figure 4A**). In the ‘digital era’, spanning the past two decades, biological data was generated and applied primarily through human-driven processes: researchers would select small numbers of sequences from public databases, analyse them using bioinformatic tools, and validate their properties through laboratory testing. In this paradigm, the limiting factors were not data availability, but rather human interpretability, experimental throughput, and the capacity of conventional computational tools (Rives et al. 2021). Most research pipelines required only hundreds to thousands of sequences, and public repositories like UniRef, GenBank, and MGnify, containing hundreds of millions of entries, were more than sufficient. The emergence of large scale models has prompted a transition to the ‘AI era’, fundamentally changing data needs as models trained to learn generalisable patterns across the full landscape of life require hundreds of millions or even billions of sequences. These are the requirements that the existing data infrastructure is increasingly unable to meet (Hwang and Cornman 2025).

The consequence is a growing mismatch: while model architectures and compute capabilities continue to scale, the available data to train on has plateaued. This shift marks the arrival of the biological data wall - a systemic bottleneck where data, not algorithms, becomes the primary constraint on progress. Without new approaches to data generation, access, and curation, biological AI will remain trapped within the narrow window of sequences supported by current public data infrastructure.

### 3.2 Uneven Origins: Where Does Biological Sequence Data Come From?

Most publicly available data comes from three domains: clinical settings (e.g., human genomes, microbiomes, pathogens), laboratories (model organisms and synthetic systems), and the natural environment. Of these, nature is by far the richest reservoir of phylogenetic and functional diversity, particularly among microbes and non-model eukaryotes, but remains the most under-sampled and poorly characterized (L. R. Thompson et al. 2017). In contrast, clinical and laboratory sources dominate existing databases due to concentrated infrastructure, routine sequencing, and clearer legal frameworks, leading to an overwhelming focus on humans, pathogens, and a few model species. This imbalance reflects longstanding disparities in scientific investment and access, and now poses a fundamental constraint on the future of biological Digital Sequence Information (DSI) (European Commission JRC 2024).

### 3.3 Sampling Bias: What and Where We Sequence Matters

The shape of today’s biological sequence databases reflects not just what exists in nature, but what has been chosen to be studied. Taxonomic and geographic biases are deeply embedded in the way biological data is generated, and these biases have direct consequences for the training and generalisation capacity of biological foundation models.

Taxonomically, public databases are dominated by a small number of species that have historically attracted the most research attention: in the SRA over 68% of all reads map to just 5 organisms **(Figure 1D).**

This is mirrored by a stark geographic bias **(Figure 4D)**, with biodiversity research disproportionately concentrated in high-income countries and regions near major institutions, leaving many species-rich areas underrepresented in the global scientific record (Tydecks et al. 2018). Tropical regions that are home to much of Earth’s endemic biodiversity such as the Amazon, Congo Basin, and Southeast Asia remain vastly underrepresented in public databases. These gaps are most often due to historical inequities in funding, infrastructure, and accessibility (Makhalanyane et al. 2023; Chavarría 2025).

### 3.4 Infrastructure and Access to Global Biodiversity Are Unevenly Distributed

Over the past decade, technological advances have dramatically lowered the cost and complexity of sequencing. Portable devices such as Oxford Nanopore’s MinION (Jain et al. 2016) have enabled field-based DNA extraction and real-time sequencing, and cloud-based platforms have made bioinformatic analysis more accessible than ever. The team at Basecamp Research was one of the first to demonstrate *in situ* genomic analysis in resource-limited settings, becoming the first to conduct fully off-grid DNA sequencing in polar environments (Gowers et al. 2019).

Meanwhile, the cost of DNA sequencing has decreased from over $10,000 per human genome in 2011 (Kris A. Wetterstrand 2019) to under $200 today (LeMieux 2025) and compute costs for training and deploying models have similarly declined, aided by more efficient architectures and commodity cloud infrastructure.

However, while the tools now exist to collect and analyse biological data at global scale, the systems needed to deploy them universally and equitably remain underdeveloped. In biodiversity-rich but resource-limited regions, sequencing infrastructure is often absent or patchy. Even where technology is available, a lack of cold-chain logistics, consistent reagent supply, or local bioinformatics expertise often limits its use (Chavarría 2025; Makhalanyane et al. 2023; Tydecks et al. 2018). Training local researchers, building stable data pipelines, and supporting long-term storage and access are still rare investments, despite being prerequisites for meaningful scientific output.

Importantly, access also depends on governance, funding, and trust. Many research institutions remain hesitant to invest in long-term infrastructure in regions with unclear legal frameworks, limited institutional continuity, or risk of regulatory backlash (S. Laird et al. 2020). As a result, tools that are technically available and increasingly affordable are often underutilised where they are needed most (Okune et al. 2018).

This disconnect has become a key contributor to the global data wall. The world’s capacity to sequence and compute is no longer the bottleneck. The bottleneck lies in our systems for deploying that capacity globally (Okune et al. 2018). Until sequencing capacity, governance infrastructure, and global incentives are better aligned, biological foundation models will remain constrained by a data landscape shaped less by what exists in nature and more by where systems happen to function.

### 3.5 Legal and Regulatory Friction: From the CBD to DSI Gridlock

Even as sequencing technology and data processing have advanced rapidly, regulatory uncertainty has emerged as one of the most significant barriers to the expansion of public biological sequence databases **(Figure 4A–C)**. At the centre of this challenge is the Convention on Biological Diversity (CBD) and its implementation through the Nagoya Protocol, which came into force in 2014 and has now been ratified by 142 countries worldwide, with over 300 separate legislative, administrative or policy measures created **(Figure 4B)** (Secretariat of the CBD 2025). While foundational for promoting conservation and ethical use of biodiversity, these frameworks have also introduced complex new layers of reputational risk and legal ambiguity, particularly for digital sequence data **(Figure 4C).**

As the authors have explained before (Vince, Gowers, and McGibbon 2024), the cumulative effect of this uncertainty (**Figure 4C**) has been to slow commercial biodiscovery and this complex regulatory environment has profound implications for the commercial application of AI in biology (**Figure 4A**). As the biotechnology sector has moved from the ‘Digital Era’ to the ‘AI Era’, the demand for data has soared, but, as the primary source of biological data is still the public online collections, the databases have not kept pace with demand. Commercial actors, wary of the legal complexity and transaction costs associated with formal bioprospecting, have shifted overwhelmingly toward open-access databases such as GenBank, UniProt, and EMBL-EBI, an approach which has been facilitated by advances in DNA sequencing and synthesis over the same time period (Halewood 2024).

A central point of contention today is the treatment of Digital Sequence Information (DSI) - the genetic data derived from physical samples. The ease with which DSI can be digitally stored, shared, and utilised across borders has underscored unresolved questions about ownership, traceability, and fair compensation, particularly for provider countries (Klünker and Richter 2022; Akpoviri, Baharum, and Zainol 2023). As a result, the debate over how to govern DSI remains one of the most polarising and legally ambiguous challenges within international biodiversity policy frameworks.

DSI is increasingly framed as a new frontier of ‘biopiracy’ - a domain in which value is extracted from biodiversity-rich regions through data, rather than samples, and redistributed without consent or benefit-sharing (Bond and Scott 2020).

Although DSI was not explicitly included in the original Nagoya Protocol, many countries have moved to regulate it under national Access and Benefit-Sharing (ABS) laws, particularly where it is used in commercial contexts (Secretariat of the CBD 2025) (**Figure 4C**, second panel). This includes some of the world’s most populous and biodiverse nations, including Brazil, India, Indonesia, South Africa, and China, which now require prior consent and legal agreements to be negotiated, not just for physical genetic samples, but for downstream digital biological sequence data (DSI) derived from them (Bagley et al. 2020). These divergent approaches have introduced complex new layers of legal ambiguity and reputational risk for global actors navigating ABS compliance, especially in cases where domestic regulations go beyond what is internationally harmonised.

Recent international efforts to address these regulatory complexities surrounding DSI culminated in the 16th Conference of the Parties (COP16) to the Convention on Biological Diversity (CBD) in 2024 Cali, Colombia. During this meeting, Parties adopted Decision 16/2, which endorsed the development of a global multilateral benefit-sharing mechanism - the ‘Cali Fund’ - specifically for DSI.

While politically significant, the COP16 Decision 16/2 leaves many questions unresolved and makes the commercial use of public biological sequence databases even more complex (Van Vooren and Gevrenova 2025). This is particularly so because the ‘Cali fund’ excludes biological sequence data uploaded to the public databases in breach of national law under the Nagoya Protocol.

The intersection of intellectual property law and ABS obligations is also becoming increasingly significant in the governance of genetic resources. In May 2024, WIPO member states adopted a new international legal measure requiring patent applicants to disclose the origin or source of genetic resources and associated traditional knowledge used in their inventions (WIPO 2024). This marks a step toward aligning the global IP framework with the principles of the CBD and Nagoya Protocol. Disclosure requirements may also apply to digital representations of biological material, including sequence data. For users of open-access databases, particularly in jurisdictions with broad ABS rules, this introduces new due diligence requirements that complicate regulatory compliance and patent protection. The challenge is especially acute for those relying on sequence data without clear provenance.

As national interpretations of digital sequence information (DSI) obligations diverge, users face growing uncertainty over compliance and potential retrospective liability, discouraging investment and hindering equitable global participation in digital biodiversity research (Bond and Scott 2020).

As in other fields, the use of training data from public repositories with disputed rights is becoming a focus of legal scrutiny and emerging regulation, introducing friction just as biological AI models increasingly depend on scale. Legal uncertainty around data provenance is not only raising the risk of litigation but also constraining the availability of the very data on which continued progress in research, innovation, and conservation depends. These dynamics slow scientific discovery and undermine global efforts to build inclusive and equitable bioeconomies (Arita 2025; Akpoviri, Baharum, and Zainol 2023).

### 3.6 Incentive Mismatches: A Disconnected Global Innovation System

At the heart of the biological data wall lies a deeper structural issue of misaligned incentives. The majority of the world’s unexplored biological diversity is concentrated in countries across the global South (Jenkins, Pimm, and Joppa 2013), while the technical capacity to transform that diversity into scientific and commercial value resides primarily in the global North (Hoffman 2006; Linck and Cadena 2024). This disconnect shapes what data is collected and whether it is collected at all **(Figure 5A, top panel)**.

This has led many biodiversity-rich countries to enact increasingly strict access regulations, slowing or even halting the flow of new data into global repositories. At the same time, private companies, concerned about legal risk, enforcement uncertainty, and the possibility of retroactive claims, often avoid working in jurisdictions with complex or ambiguous access and benefit-sharing rules.

The result is a negative feedback loop. Limited benefits reduce local incentives to invest in sequencing, curation, or data sharing. Regulatory friction deters external investment and collaboration (Halewood 2024). Public repositories, required to operate under open-access principles, lack mechanisms to ensure equitable benefit-sharing (S. Laird et al. 2020), further discouraging contributions from countries with high biodiversity but limited leverage. Over time, this has hardened into a structural asymmetry: those with the most biodiversity contribute the least data, and those with the most data infrastructure control most downstream applications. For biological AI, this has direct consequences: models are trained on data that reflect historical convenience, not global biological reality **(Figure 1D and 4D).**

## 4. Basecamp Research’s Data Supply Chain: A Scalable and Incentivised Model for Ethical and Equitable Data Generation

### 4.1 Reimagining the Supply Chain: From Passive Collection to Active Partnership

The traditional model of biological sequence data generation has been largely passive and academically driven. Public databases such as UniProt have accumulated data over decades, sourced from more than 400,000 contributors across varied research institutions and studies (“UniProt” 2025). While this decentralised approach supports openness and reproducibility, it also imposes limits on data scale and quality, and is often accompanied by fragmented or ambiguous regulatory frameworks, particularly concerning commercial use **(Figures 2B–C, 4C)**. Importantly, as public databases operate as an airgap between the commercial data used in biotechnology and the data source in biodiversity, benefit-sharing with source countries or communities is almost nonexistent (Halewood et al. 2025) **(Figure 5A, *top panel*)**.

Basecamp Research set out to build a fundamentally different kind of supply chain that treats biodiversity providers as active partners (**Figure 5A**, ***bottom panel***). Since 2019, Basecamp has developed and operated a scalable, partnership-based data infrastructure that spans 26 countries and autonomous regions to date. At its core is a commitment to pre-permissioned commercial use: no data is ingested into the database unless terms for commercial access have already been negotiated with the relevant stakeholders, including national authorities (usually the Competent National Authority), landowners, and local institutions (**Figure 5A**-**C**).

This model aligns the monetary and non monetary incentives of all parties. For commercial users, it ensures that any data accessed via BaseData can be deployed immediately without the delays, risk, or ambiguity that typically accompany commercialisation of open-access data. For biodiversity providers, it ensures that data usage is traceable, reportable, and contractually governed, with both monetary and non-monetary benefits flowing back as data is used in commercial R&D. This approach enables rapid, scalable growth while embedding equity, compliance and accountability at every step.

### 4.2 Embedding Compliance: Built-In Permissions and Traceability

At the heart of Basecamp Research’s data supply chain is a framework designed to ensure that every sequence is not only biologically valuable, but legally and ethically usable, particularly in high-stakes commercial contexts. Unlike public databases, which typically separate data content from its regulatory provenance, BaseData embeds access permissions and benefit-sharing terms directly into its data structure (**Figure 5B**).

This structure ensures that every piece of data is fully traceable to its source and aligned with the conditions under which it was negotiated, permitted and collected. It also enables frictionless compliance with evolving international frameworks such as the Nagoya Protocol and national regulations around the use of digital genetic data. Importantly, because commercial usage rights are embedded from the outset, BaseData can be used for biotechnology R&D, AI model training, and product development without the risk of delayed commercialisation permissions or *post hoc* negotiations.

For model builders and industrial users, this legal clarity enables faster operations, greater certainty, and improved scalability - critical prerequisites for deploying biological foundation models at scale, especially in commercial settings. For biodiversity providers, it ensures that ownership and free and informed consent are recognised and enforceable, fostering a data ecosystem that is not only scalable, but fundamentally fair.

### 4.3 Future-Proofed Benefit Sharing for the AI Era

Historically, monetisation of genetic information centred around the discovery of individual sequences that could be isolated, engineered, and expressed, typically in pharmaceuticals, agriculture, or industrial biotechnology. In these cases, benefit sharing occurred, if at all, only after years or decades of product development (Wynberg, Schroeder, and Chennells 2009; S. A. Laird and Wynberg 2018; Ijinu et al. 2022). However, in the AI era **(Figure 4A),** biological data is no longer just a means to find an individual useful molecule; as a corpus it can be used as a foundation for model training, embedding biological knowledge into systems that may produce thousands or millions of novel candidates (Henshall 2024). The data traceability built into BaseData allows the tracking of data used for AI training and is therefore able to determine commercial monetary benefits such as royalties proportional to the amount of data used from a given agreement provenance, whether the commercial product uses natural sequences, sequences designed by an AI model or a combination of the two **(Figure 5B and 5C)**. This data-driven, proportional model eliminates the ambiguity that often complicates benefit-sharing under public databases data and national ABS frameworks. This model avoids assigning value based on subjective assessments of data utility, or phase of training, and instead ties compensation directly to objective inclusion in the training dataset. It is simple to calculate, scalable across evolving AI workflows, and, critically, transparent to all stakeholders.

### 4.4 Delivering Value and Building Capacity: A New Standard for Inclusive Participation

Basecamp Research’s data supply chain is designed to generate the world’s richest biological sequence dataset and to ensure that those who contribute to it (the biodiversity custodians, stakeholders and rightholders) are included in the value it creates. This model delivers value across multiple time horizons. In the short term, it provides upfront investment in scientific capacity through financial support, hands-on training, and portable, state-of-the-art molecular biology facilities. Over the medium term, it strengthens collaboration through continued employment, data sharing and supporting publications and research priorities set by partner institutions. In the long term, the model embeds data traceability mechanisms, enabling the transparent calculation and distribution of commercial royalties derived from the data.

Unlike traditional access and benefit sharing frameworks, which often delay benefit sharing until a final product reaches market, sometimes decades after data collection (S. Laird et al. 2020), the Basecamp Research supply chain allows royalty disbursements to be triggered at the point of data use and not only at the point of final product commercialisation. By the end of 2024, Basecamp Research had shared commercial royalties with 52 beneficiaries across 19 countries, with the fastest turnaround from sample collection to payment recorded at just 9 months (**Figure 5E**). **Figure 5D** illustrates Basecamp’s broader commitment to equitable partnership development beyond royalty payments.

This ensures that biodiversity providers are not only compensated for their data but are equipped with the tools, skills, and opportunities to generate new data, apply it autonomously, and maintain an active role in the global bioeconomy. Example studies arising from our biodiversity partnerships, include one on antibiotic resistance (Clark et al. 2025) and one on seabed species monitoring (Bolton et al. 2024).

By embedding both economic and scientific value into its partnerships, Basecamp’s model sets a new standard for equitable data generation. It turns biodiversity access from a one-time transaction into a long-term, sustained shared investment that can scale with the needs of AI models and thereby strengthening national capacities, fostering inclusive innovation, and advancing the underlying objectives of the original Convention on Biological Diversity.

## 5. A New Data Supply Chain: A Global Engine for Rapid, High-Quality Biological Data Generation

### 5.1 Reaching the Unreachable: Global Sample Collection in Extreme Environments

To ensure uniform data collection from any conceivable environment we developed a suite of mobile, field-deployable molecular biology tools and protocols that enable real-time, on-site DNA extraction and analysis, without the need for large-scale laboratory infrastructure (**Figure 6A**). These tools have been deployed across an expanding global network of partner labs, enabling high-quality sequencing in some of the world’s most extreme and logistically remote environments (**Figure 6C)**.

**Figure 6:**
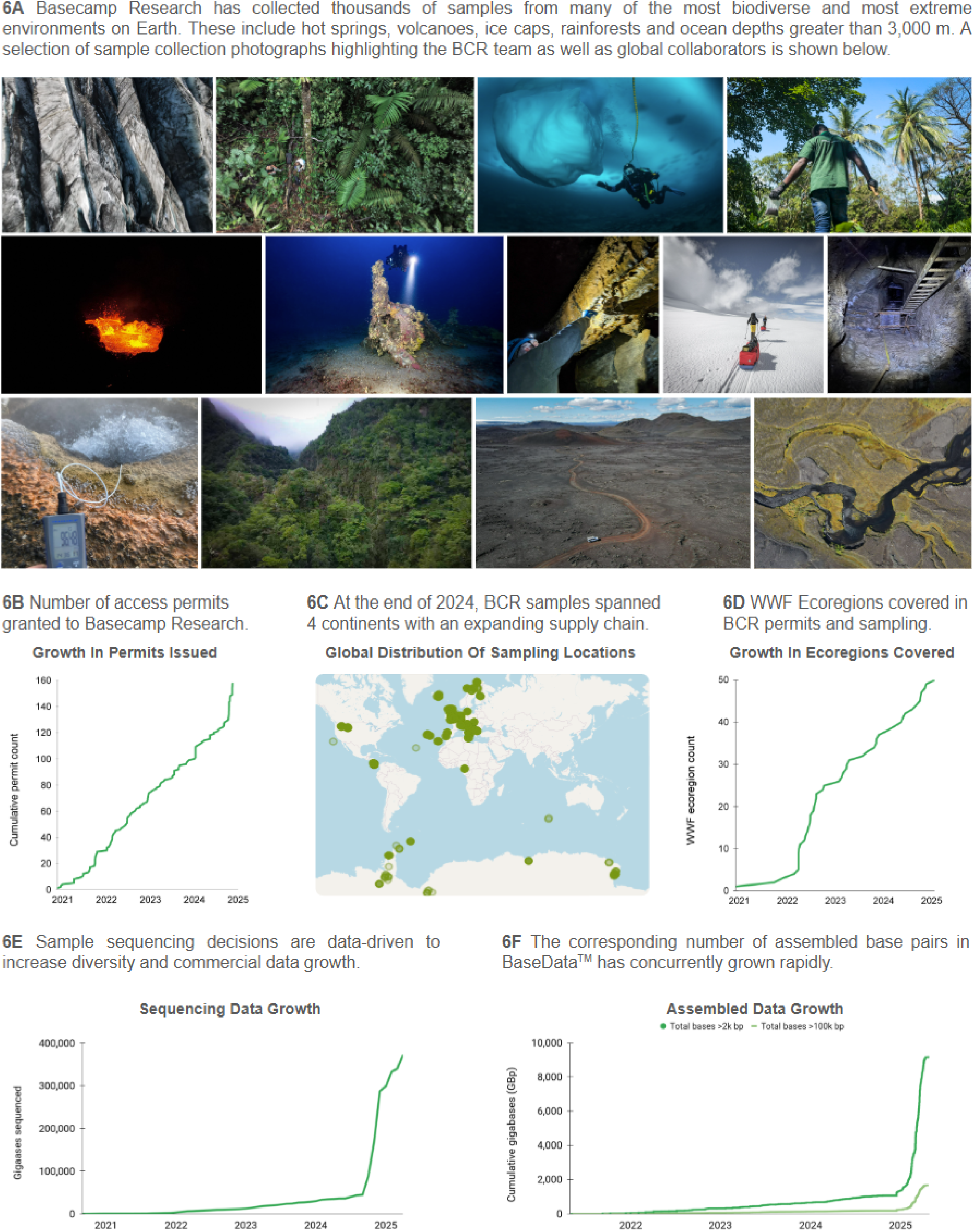
Basecamp Research’s data supply chain - rapid operational growth

Basecamp Research’s global sampling network is supported by more than 150 active commercial access permits and benefit sharing focused collaborations with protected areas such as national parks, nature reserves and other private landowners, national management areas and regulatory local authorities (**Figure 6B**). By the end of 2024, it had enabled sample collection across four continents and across 50 distinct WWF ecoregions (“Terrestrial Ecoregions of the World” 2012), with further geographic expansion already underway in 2025 (**Figures 6C–D**).

Importantly, this infrastructure allows data collection to be targeted and data-informed, driven by both prior findings and the known gaps in global sequence coverage. By designing a supply chain around novelty, accessibility, and regulatory clarity, this model moves beyond opportunistic data collection, sequencing and toward a systematic mapping of the planet’s biological potential.

### 5.2 Driving Data Growth Through Data-Driven Sampling

Basecamp Research’s global sampling infrastructure also enables data generation at scale, tuned for maximum biological and computational value. In contrast to traditional field-based sequencing efforts involved in the growth of public datasets, which often prioritise specific academic questions, the approach presented here is iterative and data-driven with a singular unified aim of optimising utility for sequence-based biological foundational model training.

Leveraging insights from previously sequenced samples to refine extraction protocols, optimise high-molecular-weight DNA preservation, and selectively target high-value biomes, we achieved a 10x increase in total sequenced bases in 2024, from under 30,000 gigabases in January 2024 to more than 300,000 gigabases by December 2024 (**Figure 6E**).

Crucially, this growth in raw sequence data translated into an even more dramatic expansion in biological insight. The number of ORFs identified grew 12-fold, from 0.8 billion to 9.8 billion over the same period (**Figure 2E**). During this growth, full traceability and compliance was maintained in alignment with Access and Benefit-Sharing requirements,ensuring that each sequence can be attributed to its source. This commitment to transparency and legal integrity creates a powerful feedback loop: as more data is generated under compliant frameworks, future collection efforts can be more precisely targeted and responsive to both scientific needs and legal obligations. In turn, this enables the dataset to be rapidly tuned to meet evolving model and policy requirements, driving growth not just in volume, but in functional utility.

### 5.3 Assembling Richer, More Useful Genomes

To improve models’ reasoning across evolutionary and functional dimensions, training sequence data must offer not just depth, but biological context: long assemblies, high gene density, intact structural features, as well as environmental and ecological metadata. The data generation pipeline presented here is designed explicitly to meet this need.

We achieve genome contiguity where over 99% of assembled bases are contained in contigs longer than 2 kilobases, with more than 18% exceeding 100 kilobases (**Figure 6F**). This is a level of resolution that allows for the capture of gene clusters, operons, regulatory elements, and mobile genetic elements within single contiguous sequences.

This structural richness is not an incidental benefit, it is foundational to enabling a new class of biological models. Long contigs make it possible to capture and learn from higher-order biological systems: the colocalisation of CRISPR arrays with Cas proteins, the integration of phage genomes into host sequences, the genomic signatures of horizontal gene transfer, and the genetic co-evolutionary patterns of protein binders, as illustrative examples. These are precisely the kinds of relationships that token-level models trained on fragmented sequences cannot resolve.

This assembly quality directly amplifies data efficiency. High-contiguity genomes reduce redundancy, enhance ORF recovery, and improve annotation accuracy, all of which increase the usable signal per base sequenced. This means that, per sample, Basecamp generates not only more data, but more learnable data, an essential requirement for training foundation models capable of generalising across the complexity of life.

## 6. Conclusion: A New Foundation for Biological AI

Foundation models are reshaping the life sciences, offering unprecedented capacity to learn, generalise, and design across the full complexity of biology. However, as this paper has shown, these capabilities are now constrained by a data wall - an infrastructure bottleneck rooted in outdated systems of biological data collection, curation, and governance. Public sequence databases, while foundational to decades of research, were not built for the demands of large-scale machine learning. Their limitations in scale, growth, redundancy, bias, fragmentation, and legal ambiguity are now emerging as primary constraints on further progress.

BaseData offers a new foundation. Purpose-built for the AI era, it combines global biological reach with regulatory clarity, ethical data sourcing, and machine learning-ready structure. By integrating benefit-sharing agreements, long-read sequencing, deep metadata, and scalable field operations, Basecamp Research has created a data ecosystem that grows not only in size, but in inclusivity, legal certainty, and scientific and commercial utility.

We propose this model as a blueprint for the next generation of biological infrastructure: one that spans ecosystems, recognises sovereign rights, accelerates global capacity, and ultimately enables AI systems to learn from, and work for, all of life on Earth. BaseData shows that a new era of biological understanding has begun, and that it needs to be built on a foundation of scale, equity, and intelligence from the start.

## 7. Extended Figure Legends

Figure 6: Basecamp Research’s Data Supply Chain - rapid operational growth

6A Basecamp Research has collected thousands of samples from many of the most biodiverse and most extreme environments on Earth. These include hot springs, volcanoes, icecaps, rainforests and ocean depths greater than 3000m. Photo descriptions and credits are below:

Top row, left to right:

1. Icecap. Photo credit: Basecamp Research
2. CENIBIOT Laboratory of CONARE collaborators and Basecamp Research team in Tirimbina Nature Reserve, Costa Rica. Photo credit: Coldhouse Collective
3. Basecamp Research team diving under ice shelf, Antarctica. Photo Credit: Joseph Marlow
4. Cameroonian collaborator taking samples in Cameroon. Photo credit: Elizabeth Dalziel

Middle row, left to right:

5. Volcano. Photo credit: Oliver Vince

6. Heritage Malta collaborator diver taking samples from WW2 shipwreck, Malta. Photo credit: Dave Gration and Heritage Malta

7. Basecamp Research team taking a biofilm swab from a cave in the Azores. Photo credit: Basecamp Research

8. Basecamp Research Co-Founders sledging the mobile lab across the Vatnajokull ice cap in Iceland. Photo credit: Basecamp Research

9. Samples taken from 51m below surface in an inactive silver mine. Photo credit: Basecamp Research

Bottom row, left to right:

10. Samples from high temperature, acidic hot springs add to the discovery of unique extremophiles. Photo credit: Basecamp Research

11. Drone footage of sampling environment from Madeira. Photo credit: Basecamp Research

12. Drone footage of the Basecamp Research portable lab on top of a jeep on the way to an icecap for sampling.

Photo credit: Basecamp Research

13. River landscape. Photo credit: Basecamp Research

## 8 Acknowledgements

The authors gratefully acknowledge the biodiversity stakeholders around the world who have granted Basecamp Research permissions to access their sites for the collection and analysis of environmental samples for metagenomic research and commercialization purposes.

The authors would also like to thank Richard De Napoli, Alexandros Papadopoulos, Neem Patel, Silvano Ulivi, Luis Enrique García-Rivera, Jonathan Finn and all other current and former employees of Basecamp Research for their contributions to this work.

We are grateful to the collaboration labs which have helped us with the collection of samples that led us to the generation of this data, including: British Antarctic Survey (collaboration led by Professor Melody Clark), Heritage Malta, Malta (Professor Timmy Gambin), CENIBiot of Conare, Costa Rica (Professor Max Chavarría and previously Professor Randall Loazia Montoya), Balaton Research Limnological Institute, Hungary (Dr. Kálmán Tapolczai), Wallowa Resources, USA (Nils Christoffersen), AJESH, Cameroon (Harrison Ajebe Nnoko), Communities of Cameroon (Ongue Yassoukou, Nkom, Ngompem, and Bibema), University of Buea (Dr. Eric Fokam), and the Scripps Oceanographic Institute, USA (Alexandra Hangsterfer).

We additionally are grateful for scientific input on this work from Professor Chris Mason, Weill Cornell Medicine and Kevin Yang, Microsoft Research. We are also incredibly grateful to the photographers who are individually credited in the Figure Legends.

## 10. Materials and Methods

### BaseData Generation

Basecamp Research operates a global data supply chain, using proprietary protocols for sampling and metadata collection from unexplored biodiversity. Prior to any sampling activity, appropriate permissions for access to genetic resources and commercial utilization of the subsequently generated data are sought from relevant stakeholders including landowners or national-level government where appropriate under national or multinational regulations.

Sampling is carried out either by the Basecamp Research in-house team of expedition scientists or by our global network of collaborator laboratories across the globe. A suite of environmental, physical, and chemical metadata is collected for each sample, and DNA is extracted using proprietary protocols. DNA is sequenced to depths targeted to maximize diversity capture using a combination of Oxford Nanopore and Illumina for long and short reads, respectively, allowing for the generation of high quality and high contiguity genomic assemblies.

A custom assembly and annotation pipeline is run for standard QC of the sequencing reads and subsequent *de novo* assembly and polishing of contigs. ORFs and non-coding RNAs are annotated on the genomic assemblies. The open reading frames are translated into amino acid sequences which are then subjected to comprehensive *in silico* annotations, including PFAM (Mistry et al. 2021), KEGG (Kanehisa Laboratories 2025), COG (NCBI 2025), InterPro (Blum et al. 2025). Functional annotations are performed with custom Hidden Markov Model-based and Deep Learning based models. Multiple custom taxonomic annotation methods were performed both on the ORF and the genomic assembly level. We also perform comprehensive multi-gene, pathway, and operon-based annotations for biosynthetic gene clusters, CRISPR arrays, and mobile genetic elements with custom Hidden Markov Model and deep learning-based models.

### BaseData and Public Genomic Uniform Manifold Approximation and Projection (UMAP)

Kmer frequencies (k=4-6) were calculated for all BaseData contigs and metagenomic contigs from OpenGenome2 using Jellyfish (2.3.1) (Marçais and Kingsford 2011). Contigs were length-filtered to include only contigs >50 kb. The data was scaled with scikit-learn standard scaler and the UMAP was calculated using *umap.UMAP* with parameters n_neighbors=15, min_dist=0.2.

### BaseData and Public Protein Uniform Manifold Approximation and Projection (UMAP)

All BaseData sequences deriving from contigs >2kb were clustered with MMseqs2 (version 15.6f452) (Steinegger and Söding 2017). This was a multi-step process due to limitations in MMseqs2 (version 15.6f452) of a maximum of 4 billion sequences. All sequences were split into six evenly sampled buckets and clustered in two steps, initially at 90% identity and 90% coverage using coverage mode 2 (to remove fragments), and then at 50% identity and 90% coverage. This method is adapted from the method used by OMG for the OMG_PROT50 (Cornman et al. 2024). Finally cluster representatives were concatenated with the singleton sequences. The sequence set was then subsampled by the same proportion as outlined in BaseData Final Database size to account for redundancy amongst buckets. The resulting set was then randomly sampled to 0.5% and embeddings calculated using ESM 650M.

Public databases (NCBI, UniProt - both Swissprot and TrEMBL) were clustered in a single step, still retaining the 2 passes (90% identity and 90% coverage, coverage mode 2, then, 50% identity and 90% coverage). A random sample of 0.5% was taken of each database and visualised using ESM embeddings and a UMAP.

The OMG_prot50 database was used as it was available, processing methods described in (Cornman et al. 2024). It was subjected to a random sampling of 0.5%, visualized with ESM embedding and a UMAP.

The inclusion or exclusion of singletons is an important topic to address: these could represent novel proteins important to include for representing as broad as possible a fitness map of protein diversity, or conversely could represent artifacts such as truncated proteins or sequencing or annotation errors. We ran extensive analysis of our singletons to gain confidence in their biological value. Results included: BaseData singletons came from long contigs (>2kb) with a high coding density (>65%), had high kmer complexity, and were made up of <50% low complexity regions. Protein fold predictions showed a significant fraction of singletons with high-confidence structures as measured by pLDDT and pTM scores. Additionally, we have expressed and tested proteins from our singleton clusters in house, so have *in vitro* confidence that there is valuable, functional, and rare biological diversity in these sequences. We applied stringent quality filters (as described above, adapted from Profluent’s methodology for PPA-1) to the BaseData singleton population to ensure high confidence for biological validity in all singletons included. In this UMAP we have presented the OMG data following the authors’ own data curation methods, which excludes all singletons. We note that due to our sequencing pipeline (which emphasizes finding biological novelty, enables longer contigs through long reads, and ensures high per-base confidence through stringent thresholds for polishing coverage) we anticipate that the BaseData singleton population likely is more highly biased towards biological novelty and away from artifacts than the OMG singleton population may be, which may be a contributing rationale for their differing curation decision.

### BaseData Final Database Size

The final BaseData database size was determined by taking the final BaseData cluster representatives from the BaseData UMAP dataset, and concatenating with it all of the singleton sequences from each BaseData exclusive bucket. The following sequence metrics and filters - a more stringent adaptation of the filtering criteria implemented in PPA-1 (Bhatnagar et al. 2025) - were applied:

● Removal of simple kmer repeats
● <50% of low complexity regions, using TANTAN (version 51)
● ORF length <8,000 amino acids
● Coding density of underlying contig >65%
● Contig length >2kb

Finally, in order to account for singleton sequences that could have clustered with each other, or with the final representatives from other buckets (ie. how much cross bucket redundancy there is), a random 10 million ORFs were taken from the singleton set and searched against the full set of singletons (from all buckets), and the full set of cluster representatives from the final pass. This was done using MMseqs2 search with the “faster” option. Subsequently the output results were filtered by percentage identity (50%) and coverage (90%).

This proportion was applied to the total set of singleton sequences, projecting the estimate for total non-redundant singletons. This projection was combined with the total of cluster representatives to give the total final number of 2.0B non-redundant clusters at 50%.

### Protein-level Diversity Comparison Between BaseData and the Open MetaGenomic Corpus Database

Sequence diversities of different classes of proteins were compared between the BaseData and the OMG database (Cornman et al. 2024), a publicly available resource of metagenomic data containing over 3 billion protein sequences. Protein classes that were compared between the two databases are listed in **Supplementary Tables ST1 and ST2.** For each protein class, Hidden-Markov Models (HMM)-based profiles of respective Pfam annotations were obtained, and hmmsearch from HMMER (v3.3.2) (“HMMER” 2025) was run with “--cut_ga” option to search for significant potential candidates from both databases surpassing domain-specific thresholds. To generate visualisations in Figure 3A-D, sequences from each of BaseData and OMG were clustered down in linear time using MMseqs2 via alignment mode 3 (i.e. Smith-Waterman based alignment). Cluster representatives were randomly sub-selected from each database, and a sequence alignment was generated using MAFFT (v7.490), and tree file was generated using FastTree (v2.1.11) for phylogenetic visualisation.

### Species-level Operational Taxonomic Units (OTU) Comparison Between BaseData and the GTDB and OMG Databases

Universal single-copy taxonomic marker genes were selected from Parks et al. (2018). For each marker gene, hmmsearch was used to search each database (BaseData, OMG, GTDB) using TIGRFAM HMM profiles (-E 0.001 –domE 0.001 and marker-specific cut-off values), and were length-filtered to remove fragmented genes. Amino acid sequences were clustered to species-level operational taxonomic units (OTU) using MMseqs2 (version 15.6f452) cluster for each marker gene (coverage mode 2 to minimise the impact of fragmented sequences) using optimal thresholds for species delineation following methods adapted from Olm et al (2020) (Olm et al. 2020). OTUs of potential eukaryotic origin were identified and filtered using Kaiju (v1.10.1). Recombinase A was chosen for downstream visualisations as it showed the highest congruency with GTDB taxonomy. OTUs were clustered to 60% using MMseqs2 using default parameters, and 10% of the sequence representatives were randomly selected. A sequence alignment was generated using hmmalign (v3.4) and trimmed using trimal (v1.5), and a phylogenetic tree was generated using FastTreeMP (v2.1.11).

### Geographical Distribution and Organism Representation in SRA Data

Metagenomic sample metadata was obtained from the NCBI Sequence Read Archive (SRA) via Google BigQuery using the nih-sra-datastore.sra.metadata public dataset. To examine geographical distribution, the dataset was filtered to include only samples with librarysource = ’METAGENOMIC’ and limited to entries containing valid country information (excluding samples where geo_loc_name_country_calc was null or marked as ’uncalculated’).

Organism bias analysis was conducted using the sequencing volume and organism columns available within the nih-sra-datastore.sra.metadata public dataset to assess the representation patterns of different species across the archive.

### Growth of Public Databases

The annualized growth of UniRef50 clusters over time was based on data retrieved from the UniProt Reference Clusters database (“UniProt” 2025). To compile the dataset, the total number of UniRef50 sequences was extracted from successive UniProt releases, as reported on the UniRef website.

### Comparison of Genomic Model Databases

The sizes of major genomic language model training datasets, reporting total nucleotide tokens and their composition by source was compiled as follows. Data for OpenGenome and OpenGenome2 were obtained from the Arc Institute’s public releases and preprints, which report a total of 300 billion nucleotides from prokaryotic and phage genomes for OpenGenome, and 8.8 trillion nucleotides spanning bacteria, archaea, eukarya, and bacteriophage for OpenGenome2 (Nguyen, Poli, Durrant, Kang, et al. 2024; Brixi et al. 2025). The OMG dataset (Open MetaGenomic corpus) was included as a representative large-scale metagenomic resource, comprising 3.1 trillion base pairs from metagenomic sources and 3.3 billion protein-coding sequences, curated from JGI’s IMG and EMBL’s MGnify repositories with stringent quality filtering and deduplication as described in the original publication and blog (Hwang 2024; Cornman et al. 2024).

### Comparison of Protein Model Databases

The sizes of protein language model training datasets, reporting both total protein sequences and the number of clustered representatives at 50% sequence identity were compiled where available. UniRef50 clusters were used as the reference for 50% identity clustering (“UniProt” 2025). The UniMeta-200B dataset comprises 0.9 billion metagenomic protein sequences as reported in recent preprints (Cheng et al. 2024). The Big Fantastic Database (BFD) contains 2.2 billion protein sequences, with 566 million clustered at 30% identity (Jumper et al. 2021). ESM3 was trained on 3.1 billion protein sequences, as described in the ESM3 preprint (Hayes et al. 2025), while Profluent’s PPA-1 (Protein Atlas v1) contains 3.4 billion full-length protein sequences, with 183 million clustered at 50% identity (Bhatnagar et al. 2025). The OMG dataset includes 3.3 billion protein-coding sequences, with a clustered subset (OMG_prot50) at 50% identity, comprising 207 million representatives (Cornman et al. 2024).

Annualized data growth at EMBL-EBI from 2017 to 2024 was generated using publicly available data from the institute’s Annual Highlights reports, accessible via the (European Bioinformatics Institute 2025). These reports provide yearly summaries of data volumes across EMBL-EBI’s major archival resources, including the European Nucleotide Archive (ENA), PRIDE, MetaboLights, and the European Genome-phenome Archive (EGA).

### Policy Analysis

The number of national laws governing access to genetic resources (Figure 4B) was captured and plot adapted from (Michiels et al. 2021). Lists of countries with ABS regulations (Figure 4C) were captured from the Analysis and Mapping of Policies of Emerging Infectious Diseases (AMP EID) resource out of Georgetown University (Centre for Global Health, Science and Security, Georgetown University 2024).

### Comparison of Patented Sequences to Public Data

Sequence data from four major patent offices (USTPO, EPO, JPO, KIPO) were retrieved from EMBL-EBI (EMBL-EBI 2025) and the complete RefSeq database (Goldfarb et al. 2025) was obtained from NCBI. A comparative analysis was performed by clustering sequences from the patent database against RefSeq sequences using MMseqs2 (Steinegger and Söding 2017). Clustering was conducted with a minimum sequence identity threshold of 90% and default coverage parameters.

Patent sequence headers belonging to “mixed clusters” (i.e., clusters containing both patent and RefSeq member sequences) were isolated. Regular expression-based pattern matching was employed to parse these FASTA headers and extract normalized, unique patent document numbers. This process yielded 182,875 unique patent documents with at least one sequence present in a mixed cluster.

To determine the denominator, all FASTA headers from the entirety of the initially retrieved patent sequence database were analysed. This identified a total of 274,571 unique patent documents within this dataset. The percentage of patent documents from the database having at least one sequence meeting the similarity criteria with RefSeq was then calculated based on these two counts.

---

**Table ST1.**
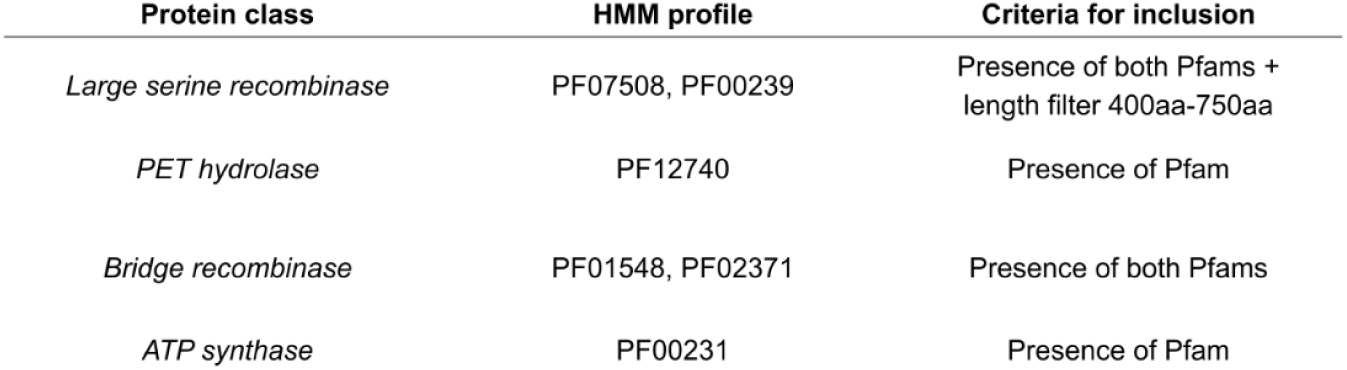
Protein Class Inclusion Criteria.

ST1 Protein Class Inclusion Criteria
| Protein class | HMM profile | Criteria for inclusion |
| --- | --- | --- |
| Large serine recombinase | PF07508, PF00239 | Presence of both Pfams + length filter 400aa-750aa |
| PET hydrolase | PF12740 | Presence of Pfam |
| Bridge recombinase | PF01548, PF02371 | Presence of both Pfams |
| ATP synthase | PF00231 | Presence of Pfam |

**Table ST2.** Sequence Clustering and Subsampling Statistics.

ST2 Sequence Clustering and Subsampling Statistics
| Count (before clustering, unique sequence IDs) |  | Sequence clustering threshold | Count after clustering |  | % Retention from original count after clustering |  | Random subsampling for phylogenetic trees |
| --- | --- | --- | --- | --- | --- | --- | --- |
| OMG | BaseData™ | (%) | OMG | BaseData™ | OMG | BaseData™ | (%) |
| 1,455,468 | 2,088,908 | 20 | 61,091 | 360,052 | 4.2 | 17.2 | 1 |
| 97,717 | 320,392 | 90 | 27,097 | 177,070 | 27.7 | 55.3 | 1 |
| 483,571 | 6,480,620 | 50 | 34,939 | 1,248,310 | 7.2 | 19.3 | 1 |
| 888,570 | 1,878,106 | 50 | 29,377 | 618,453 | 3.3 | 32.9 | 1 |

**Figure S1:**
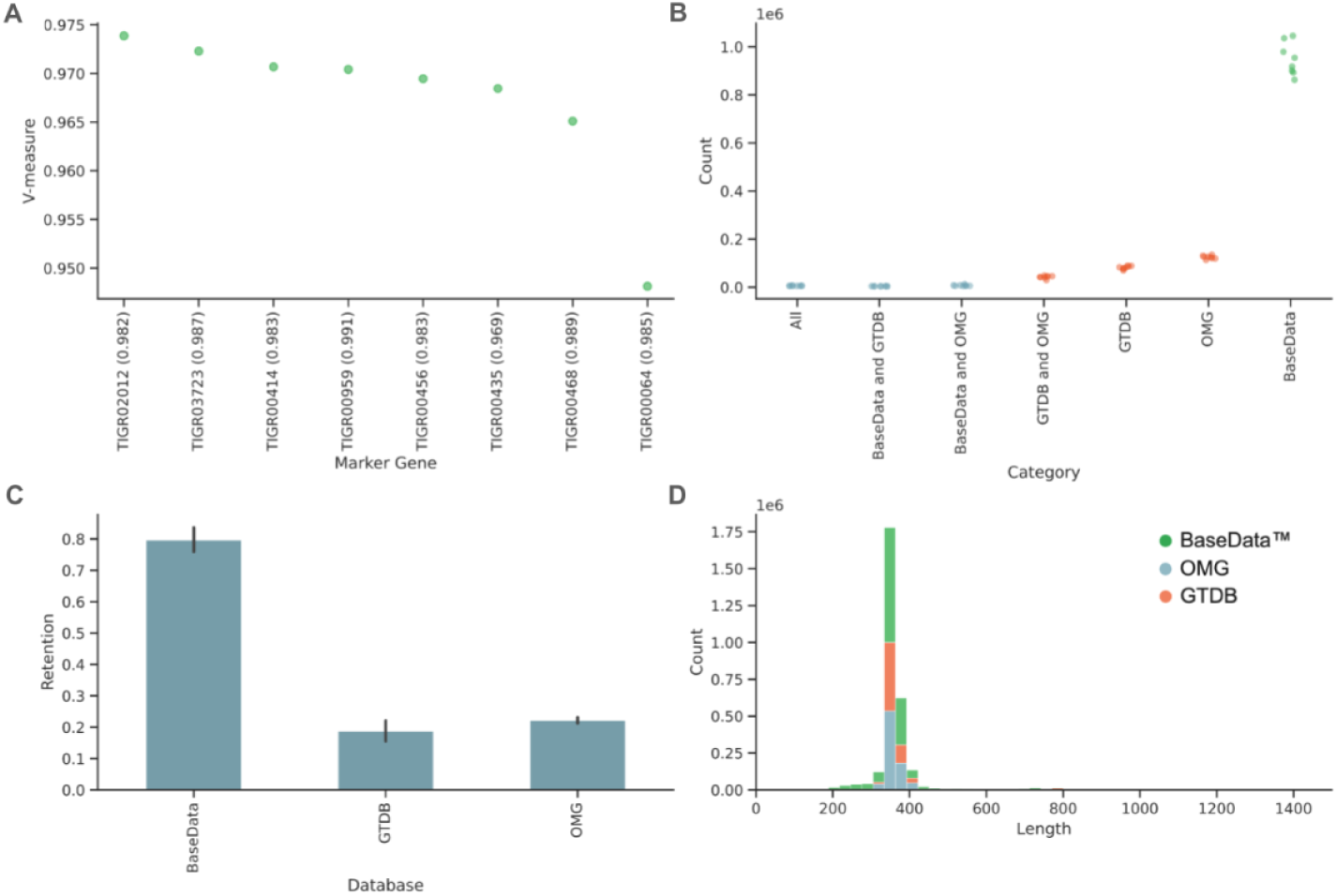
S1A (A) Accuracy of OTU clustering expressed as V-measure (harmonic mean of homogeneity and completeness) for the highest-ranking marker genes (V-measure > 0.948); clustering thresholds used are indicated in brackets. (B) Cluster counts by database overlap show varying degrees of exclusivity and intersection. (C) Clustering retention (%) of species-level OTUs across databases. (D) Length distribution of the RecA protein (the top-ranked marker gene) across BaseData™, OMG, and GTDB databases.

